# Development of a protein synthesis network at the sarco/endoplasmic reticulum in adult cardiac myocytes

**DOI:** 10.1101/2024.09.20.614189

**Authors:** Christoph Sandmann, Viviana Sramek, Sören Brandenburg, Dennis Uhlenkamp, Marc Renz, Bianca Meyer, Clara Sandmann, Nicole Herzog, Julia Groß, Hugo A. Katus, Stephan E. Lehnart, Mirko Völkers, Shirin Doroudgar

## Abstract

**Introduction:** The endoplasmic reticulum (ER) is the site of synthesis and folding of membrane and secretory proteins, which constitute a large fraction of the total protein output of a mammalian cell. Striated muscle cells contain a specialized membrane system known as the sarcoplasmic reticulum (SR) that controls calcium homeoastasis and contraction, however the biochemical and physiological relationship between the ER and SR and how both compartments participate in protein synthesis remains incompletely understood.

**Methods:** Protein quantification and imaging of selected marker proteins for ER and SR functions was performed to characterize the relationship of the ER and SR and its individual involvement in protein synthesis in neonatal and adult cardiac myocytes. Superresolution microscopy was used to examine the interaction of ribosomes and the SR in adult cardiac myocytes.

**Results:** Quantification of ER and SR-associated proteins of isolated ventricular cardiac myocytes showed that relative expression of ER/SR resident protein quality elements, as well as relative ribosome levels are decreased in adult cells, whereas SR-associated Ca^2+^ handling proteins increase. Immunocytoflourescence revealed that the membrane compartment that exists in early postpartum resembles mostly the ER and decreases in postnatal development. The SR is the main membrane network that exists in the adult cardiac myocytes, replacing the ER, in all but the perinuclear region. Immunocytoflourescence staining further indicated that both networks perform overlapping but distinct, specialized functions, such as localization of excitation-contraction coupling exclusively to the SR or initiation of secretion via the classical secretory pathway mainly from the ER. Ribosomes and mRNA were localized both in close proximity to the ER and the SR of adult ventricular cardiac myocytes. Superresolution microscopy confirmed that both the ER as well as the developed SR bind ribosomes and are direct sites of protein synthesis and protein homeostasis in adult cardiac myocytes.

**Conclusion:** Our findings suggest molecular differentiation and structural organization of the ER/SR in cardiac muscle development, resulting in the development of a protein synthesis network at the sarco/endoplasmic reticulum in adult cardiac myocytes.

## Introduction

The endoplasmic reticulum is a membrane network that when studded with ribosomes (known as the rough ER; rER), is the major site of secreted and membrane protein synthesis in many eukaryotic cell types [1]. Nevertheless, this essential organelle remains largely uncharacterized in cardiac muscle cells (myocytes). In striated muscle cells, including cardiac myocytes, a network of membranes called the sarcoplasmic reticulum (SR) has been defined and studied with important roles in muscle contraction [2]. Biochemical and structural studies of developing muscle have shown that the development of the SR is the result of the specialization of the internal membrane system [3-6]. Such a development has been proposed to arise from the differentiation of the ER, resulting in the formation of the developing SR continuous with the ER [3,7,8]. Therefore, the SR is often described as a ‘specialized form’ or ‘extension’ of the (smooth) ER of striated muscle cells [9-11]. However, the relative location and protein synthetic functions of the membrane network in cardiac myocytes remains unclear, specifically if both an ‘ER’ and a ‘SR’ network co-exist in mature cardiac myocytes or if a single highly specialized ‘SR’ membrane system fullfills both functions classically associated with the ER (e.g. protein synthesis, secretion) and SR (Ca^2^ homeoastasis and excitation contraction coupling) is unknown [12,13].

The mature SR makes contacts with invaginations of the plasma membrane, known as T-tubules as well as sarcomeres, and is responsible for the storage, release, and reuptake of Ca^2+^. It is divided into two functionally and structurally distinct regions. The longitudinal SR contains few ryanodine receptors (RyR), which mediate calcium-induced calcium release, but many SR Ca^2+^ pumps (i.e., the sarco/endoplasmic reticulum Ca^2+^-ATPase; SERCA), which transport Ca^2+^ from the cytosol into the SR lumen. The junctional SR, on the other hand, contains many RyR, for the fast, localized SR Ca^2+^ release during systole, resulting in fast activation of contraction [14,15]. Although the function of the SR in cardiac contractile Ca^2+^ handling is well established, its roles in protein synthesis, folding, and quality control in cardiac myocytes are not as clear [12]. Since most Ca^2+^ handling proteins reside in either the lumen of the SR or in the SR membrane [9,12], it is reasonable to posit that contractile Ca^2+^ handling proteins in cardiac myocytes are synthesized on ribosomes associated with the SR, similar to the way that ER-luminal and transmembrane proteins in non-cardiac myocytes are synthesized on ribosomes associated with the ER. In addition, due to its close interaction with the sarcomere, ribosomes attached to the SR may localy synthesize large sarcomeric proteinance for structural homeostasis, as recently suggested for ribosomes and mRNA close to the sarcomere [16-18].

In the present study, we examined the relative locations, postnatal maturation and protein synthetic functions of the internal cellular membrane network in primary rat cardiac myocytes. We found that with cellular maturation, cardiac myocytes develop two distinct specialized membrane networks to which proteins classically associated with ER or SR functions localize differently. Importantly, we show that the membrane networks classically described as the ER and SR of mature adult cardiac myocytes are both bound by ribosomes and are associated with active mRNA translation.

## Methods

### Cultured cardiac myocytes

Isolation of neonatal rat ventricular cardiac myocytes was performed as previously described [19]. Cells were isolated from one to two-day old WISTA rats (WISTAR RjHan:WI, Janvier Labs) via enzymatic digestion and purified by Percoll density gradient centrifugation. For western blot experiments cardiac myocytes were then plated at a density of 0.5 × 10^6^ cells per well on 34.8 mm plastic plates that had been pre-treated with 5 µg/ml fibronectin (cat# F1141, Sigma-Aldrich) in serum free DMEM/F12 medium (cat# 11330032, Thermo Fisher Scientific) for one hour and then cultured in DMEM/F12 1:1, containing 10% fetal bovine serum, 100 units/mL of penicillin, 100µg/mL streptomycin and 292 µg/ml glutamine (cat# 10378016, Thermo Fisher Scientific). As the expression of several ER quality response proteins is regulated by medium serum levels, neonatal and adult cardiac myocytes were both cultured in serum free conditions. 24h after plating, neonatal cardiac myocytes were washed once with DMEM/F12 1:1 and medium was changed to DMEM/F12 1:1, containing 100 units/mL of penicillin, 100µg/mL streptomycin and 292 µg/ml glutamine for 24h after which the cells were lysed for western blotting or fixed for immunocytoflourescence as described below. Adult mouse and rat ventricular myocytes were isolated from 10-week old C57/BL6-J mouse hearts or 8-week old WISTA rats (WISTAR RjHan:WI, Janvier Labs) respectively via enzymatic digestion as previously described [20,21]. Cells were plated in Medium 199 (cat# M7528, Sigma-Aldrich) supplemented with 100 units/mL of penicillin, 100µg/mL streptomycin and 10µg/mL Laminin (cat# L2020, Sigma-Aldrich). After one hour, media was changed to M199 media supplemented with 100 units/mL of penicillin, 100µg/mL streptomycin and lysed similar to neonatal cardiac myocytes.

### Immunocytoflourescence

Cardiac myocytes were isolated as described above. Cardiac myocytes were then plated on glass chamber slides that had been pre-treated with fibronectin for one hour. The cells were washed twice in PBS and then fixed with 4% PFA for 20 minutes. Slides were washed three times for 5-10 minutes in PBS and then incubated for 10 minutes in permeabilization buffer consisting of PBS, 0.2% Triton X-100 and 0.1M Glycine. Then slides were washed once with PBS and then incubated with blocking buffer containing 10% horse serum and 0.2% Triton X-100 in PBS for one hour. Next, slides were incubated overnight at 4°C with primary antibody diluted in blocking buffer. Primary antibodies used were SERCA2 (cat# sc-8095, N-19, Santa Cruz Biotechnology; 1:200), Ryanodine Receptor (cat# ab2827, Abcam; 1:50), KDEL (cat# 10C3, ADI-SPA-827, ENZO Life Sciences; 1:50), P-S6 Ribosomal Protein^S235/236^ (cat# 2211, Cell Signaling Technology), HA-Tag (cat# C29F4, Cell Signaling Technology), GM130 (cat# 610822, BD Biosciences; 1:50), TGN38 (cat# sc-33784, M-290, Santa Cruz Biotechnology; 1:50), CD63 (cat# MCA4754GA, Bio-Rad; 1:50). On the next day slides were washed three times for 5-10 minutes in PBS and then incubated with secondary antibody in blocking buffer for one hour. Secondary antibody used were Fluorescein (FITC)-conjugated AffiniPure Donkey Anti-Mouse IgG (cat# 715-095-151, Jackson Immuno Research; 1:100), Fluorescein (FITC)-conjugated AffiniPure Bovine Anti-Goat IgG (cat# 808-095-180, Jackson Immuno Research; 1:100, Fluorescein (FITC)-conjugated AffiniPure Donkey Anti-Rabbit IgG (cat# 711-095-152, Jackson Immuno Research; 1:100), Cy3-conjugated AffiniPure Donkey Anti-Mouse IgG (cat# 715-165-151, Jackson Immuno Research; 1:100), Cy3-conjugated AffiniPure Donkey Anti-Rabbit IgG (cat# 711-165-152, Jackson Immuno Research; 1:100). The slides were then washed three times for 5-10 minutes in PBS and incubated in a 1:500 solution of 1mM TO-PRO-3 iodide (cat# T3605, Thermo Fisher Scientific) diluted in PBS for 10 minutes. Then slides were washed twice in PBS for 10 minutes and covered with VECTASHIELD Antifade Mounting Medium (cat# H-1000, Vector Laboratories) and a glass plate and visualized by microscopy. Poly(A) mRNAs was visualized using a complementary RNA probe labeled with ATTO 647NN. Images were obtained using a Leica DMi8 confocal laser scanning microscope.

### STimulated Emission Depletion (STED) microscopy

Ventricular cardiac myocytes from wild-type mice (10 weeks old, C57BL/6N background) were isolated using a *Langendorff*-perfusion setup. Mouse hearts were cannulated and perfused with fresh, nominally Ca^2+^ free isolation buffer (in mM: NaCl 120.4, KCl 14.7, KH_2_PO_4_ 0.6, Na_2_HPO_4_ 0.6, MgSO_4_ 1.2, HEPES 10, NaHCO_3_ 4.6, taurine 30, 2,3-butanedione-monoxime 10, glucose 5.5, pH 7.4 with NaOH), followed by collagenase type II (Worthington) containing isolation buffer in order to digest cardiac tissue. After plating on laminin coated coverslips, cardiac myocytes were fixed with 4% PFA for 5 min, and incubated with blocking/permeabilization buffer overnight (10 % bovine calf serum and 0.2 % Triton in PBS). Subsequently, cardiac myocytes were incubated with primary antibodies in blocking buffer at 4°C overnight as follows: P-S6 Ribosomal Protein^S235/236^1:500 (cat# 2211, Cell signaling, Danvers, MA); RyR2 1:500 (cat# MA3-916, Thermo Scientific, Waltham, MA); and PLN 1:500 (cat# ab2865, Abcam, Cambridge, England). After washing in blocking buffer, samples were incubated with secondary antibodies diluted 1:1000 for 2 h at room temperature: goat anti-rabbit STAR635P (cat# 2-0012-007-2, Abberior, Germany), and goat anti-mouse STAR580 (cat# 2-0002-005-1, Abberior, Germany). Finally, mounting medium (ProLong Gold Antifade Mountant, Thermo Fisher Scientific, Waltham, MA) was used to embed the samples after washing in PBS. For imaging, a Leica TCS SP8 STED system with a HC PL APO C2S 100x/1.40 oil immersion objective was used. Confocal images were acquired using the following parameters: pixel size: 116.34 x 116.34 nm; pixel dwell time: 400 ns; scanning speed: 600 Hz; 16x line averaging; fluorophore excitation by a white-light laser at 635 and 580 nm; and fluorescence detection between 650-700 nm and 600-630 nm, respectively. For acquisition of STED images, an additional STED depletion laser at 775 nm was used. Images were acquired at a resolution of 16.61 x 16.61 nm. For optimal signal resolution of STAR635P and STAR580, the STED laser beam was adjusted accordingly. Fiji (https://imagej.net/Fiji) was used for raw image processing, and line profiles were plotted in GraphPad Prism 7.0.

### Immunoblotting

Cultured cells were lysed in RIPA Buffer consisting of 50mM Tris pH 7.5, 150mM NaCl, 1% Triton X-100 and 1% SDS, which was supplemented with protease inhibitor cOmplete ULTRA (cat# 05892791001, Roche) phosphatase inhibitor PhosSTOP (cat# 04906837001, Roche). Lysates were cleared by centrifugation at 4 °C for 10 minutes at 20.000 rcf. Lysate protein concentration was determined using DC Protein Assay Kit II (cat# 5000112, Bio-Rad). Equivalent amounts of protein were brought up to similar volume, mixed with Laemmli Sample Buffer (Bio-Rad; 161-0747) and 2-Mercapoethanol (cat# M6250, Sigma-Aldrich) and boiled at 95°C for 5 minutes. Samples were separated on SDS-PAGE gels and transferred to Immobilon-P transfer membranes (cat# IPVH00010, Merck Millipore). The following antibodies were used to probe the membranes: Ryanodine Receptor (cat# ab2827, Abcam, 1:1000), SERCA2 (cat# sc-8095, N-19, Santa Cruz Biotechnology; 1:1000), KDEL (cat# 10C3, ADI-SPA-827, ENZO Life Sciences; 1:2,000-5,000), Phospho-Ribosomal S6 Ser235/236 (cat# D57.2.2E, 4858, Cell Signaling Technology, 1:5000) and Ribosomal S6 Ser235/236 (cat# 54D2, 2317, Cell Signaling Technology, 1:1000). Ponceau solution was prepared with Ponceau BS (cat# B6008, Sigma Aldrich).

### Statistical Analysis

Statistical analysis was performed using GraphPad Prism 7.0 (Graphpad Software Inc; www.graphpad.com). Data values are mean ± standard error of the mean (SEM). For statistical analysis the unpaired 2-tailed *t* test was used. p < 0.05 was defined as significant difference.

## Results

To examine the changes and relative localization of functions classically associated with the ER (protein synthesis and secretion) and SR (Ca^2^ homeoastasis and excitation contraction coupling) in cardiac myocyte maturation, we used primary cardiac myocytes isolated from rat neonatal (1-3 days old) and adult (8 weeks old) [22] ventricles to measure protein levels and localization of key proteins involved in the respective functions (Figure 1*A*). We used cardiac ryanodin receptor 2 (RyR2) and sarcoplasmic/endoplasmic reticulum calcium ATPase (SERCA2) as markers of Ca^2+^ homeostasis and excitation contraction coupling. The chaperones GRP78 and GRP90, as well as the protein disulfide isomerase family A member 6 (PDIA6) were used as indicators of protein processing and quality control and the phosphorylated and unphosphorylatied ribosomal subunit S6 (RPS6) as marker of ribosome quantity and activity. We found that in neonatal compared to adult cardiac myocytes, the relative expression of ER/SR resident protein quality elements, as well as relative ribosome levels are increased. Conversely, in adult cardiac myocytes, relative protein levels of Ca^2+^ channels expression is increased (Figure 1*B*). Taken together, the expression data indicate that in association with increased contractile demand a relative switch from functions classically associated with the ‘ER’ in immature cardiac myocytes towards ‘SR’ functions in adult cardiac myocytes occurs with cellular maturation.

**Figure 1.**
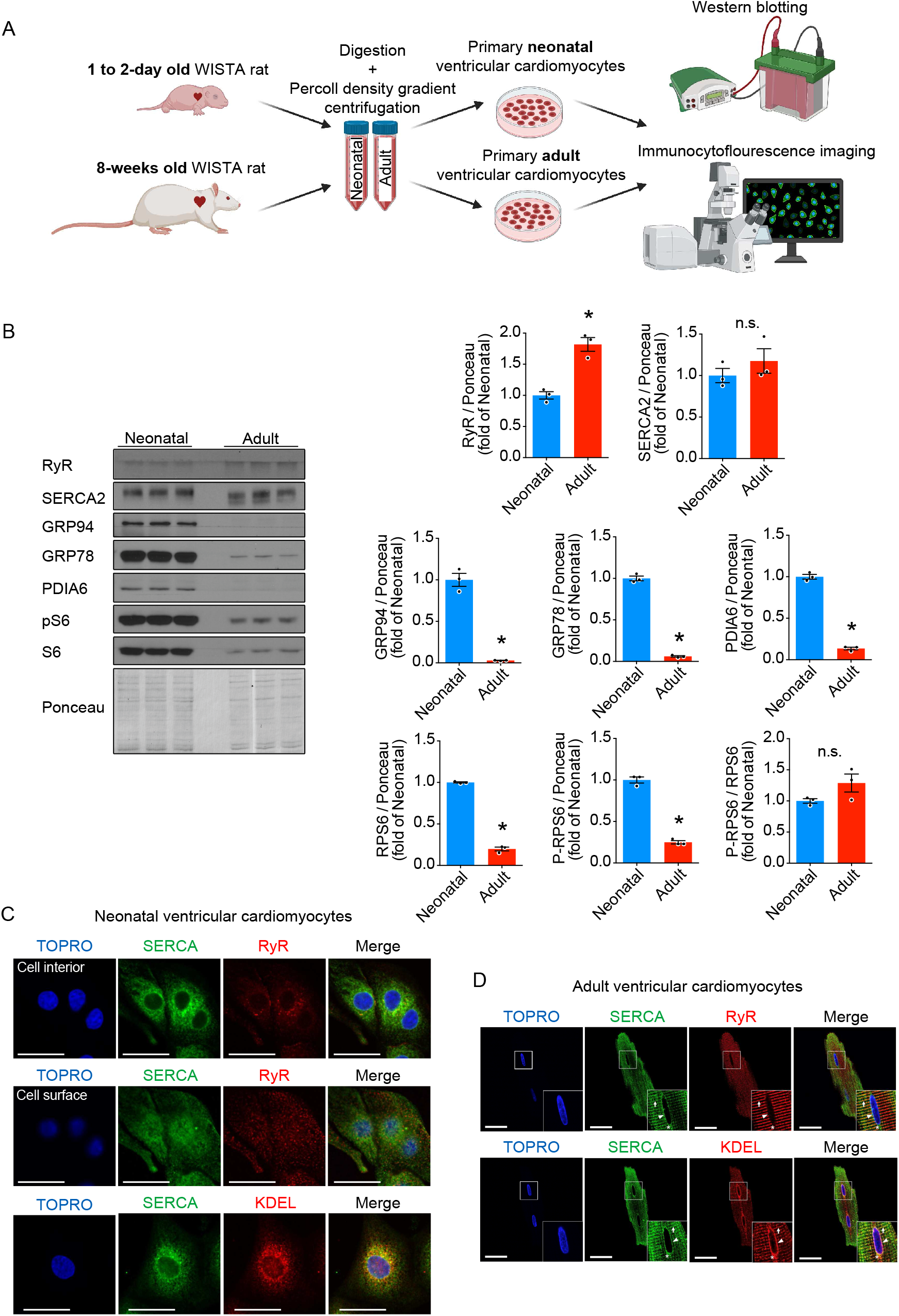
The membrane system of adult cardiac myocytes consists of two distinct networks, a perinuclear ER and a peripheral SR. **(A)** Cardiac myocytes were isolated from neonatal and adult rat hearts and subjected to immunoblotting and immunocytoflourescence. **(B)** Immunoblots and respective quantification of neonatal and adult rat ventricular cardiac myocytes for markers of Ca^2+^ homeostasis and excitation contraction coupling (RyR2 and SERCA2), protein processing and quality control (GRP94, GRP78 and PDIA6) and ribosome quantity and activity (RPS6 and p-RPS6^S235/236^). **(C)** Coimmunofluorescence labeling of neonatal rat ventricular myocytes with antibodies to the Ca2+-ATPase of the SR (SERCA) and either the cardiac ryanodine receptor (RyR2) or an antibody that specifically recognizes proteins containing the KDEL sequence, used as an ER marker. **(D)** Coimmunofluorescence labeling of adult rat ventricular myocytes as described in (D). TOPRO, nuclei; bars, 20 µm;* = p < 0.05 from neonate.

### The sarcoplasmic and endoplasmic reticulum of cardiac myocytes are interconnected compartments and have different but overlapping functions

To distinguish between ER and SR functions of the cardiac myocyte membrane network, we performed immunofluorescence labeling of excitation contraction coupling membrane proteins in neonatal and adult rat ventricular myocytes for RyR2, SERCA2, and the tetrapeptide KDEL, which marks proteins that are retained in the ER in other cell types [23]. In neonatal cardiac myocytes, SERCA2 diffusely labeled a peri-nuclear membrane network, similar to the appearance of the ER in other cell types (Figure 1C). In contrast, RyR2 was found in small peri-nuclear clusters and more evident in sharp dotted structures when the focus was shifted upwards from the nucleus, likely representing the cell surface (Figure 1C), which is in agreement with previous findings showing that RyR2 is mostly localized near the cell membrane in immature cardiac myocyters and from there travels closer to the interior of the cell during maturation [24]. Differences of SERA2 and RyR2 localitation may be indicative of the developing membrane network, and represents sites of forming junctions with the sarcolemma, known as T-tubules. KDEL staining in neonatal cardiac myocytes colocalized precisely with SERCA2 (Figure 1C), consistent with an intersection of protein homeostasis and Ca^2+^ homeostasis in a peri-nucelar membrane network that resembles the ER of other non-muscle cells. In adult cardiac myocytes, two distinct membrane networks were observed, which appeared interconnected by confocal microscopy, comprising of a peri-nuclear ER-like network with overlapping SERCA and KDEL staining that was connected to the nuclear envelope, and a peripheral striated SR, with overlapping SERCA, RyR2 and weaker KDEL staining (Figure 1D). Even though individual functions of those membrane networks seem to be overlapping and they appear interconnected by confocal microscopy, for simplicity we use the terms ER for the peri-nuclear membrane network and SR for the striated membrane network in this manuscript. In adult cardiac myocytes RyR2 is not localized in the nuclear envelope (marked by arrowhead) and not to an ER-like peri-nuclear membrane network (marked by* and KDEL staining), but is mainly localized to the SR (marked by arrow) (Figure 1D). As such, RyR2 may be used to distinguish between the ER and SR in adult cardiac myocytes. In contrast, KDEL, which is used as an ER marker in other cell types, does not localize exclusively to the peri-nuclear ER-like membrane network of adult cardiac myocytes but can also be found in the striated SR.

The ER of non-myocytes is the site from which protein secretion of the classical secretion pathway is initiated [25]. In the classical secretory pathway, proteins are synthesized on ER-bound ribosomes and translocated into the ER, transported to the Golgi and subsequently secreted after secretory vesicles fuse with the plasma membrane [26]. To further characterize the individual functions of the ER and SR of cardiac myocytes we studied whether both membrane networks are associated with classical protein processing for secretion, indicate of functions classically associated with the ER of other cell types. First, we mapped the relative localization of the classical secretory pathway to the ER and SR of neonatal and adult rat ventricular myocytes. Both the cis-Golgi marker GM130, as well as the trans-Golgi marker TGN38, were found in close proximity to the peri-nuclear ER sub-compartment in neonatal and adult cardiac myocytes (Figure 2A and B). Staining for CD63, marking late endosomes and related secretory organelles [27], however, was distributed throughout the cell, suggesting that endosomes may travel freely throughout the cardiac myocyte after being formed at the ER-associated Golgi (Figure 2 A and B). Together, our data suggest the existence of a peri-nuclear ER and a peripheral SR network in adult differentiated cardiac myocytes with specialized but overlapping functions. While protein secretion of the classical secretory pathway is initiated primarily from the peri-nuclear ER and excitation contraction coupling is located to the peripheral SR, both membrane networks appear to share overlapping functions as indicated by the localization of some membrane proteins and chaperones to both compartments [20,28,29].

**Figure 2.**
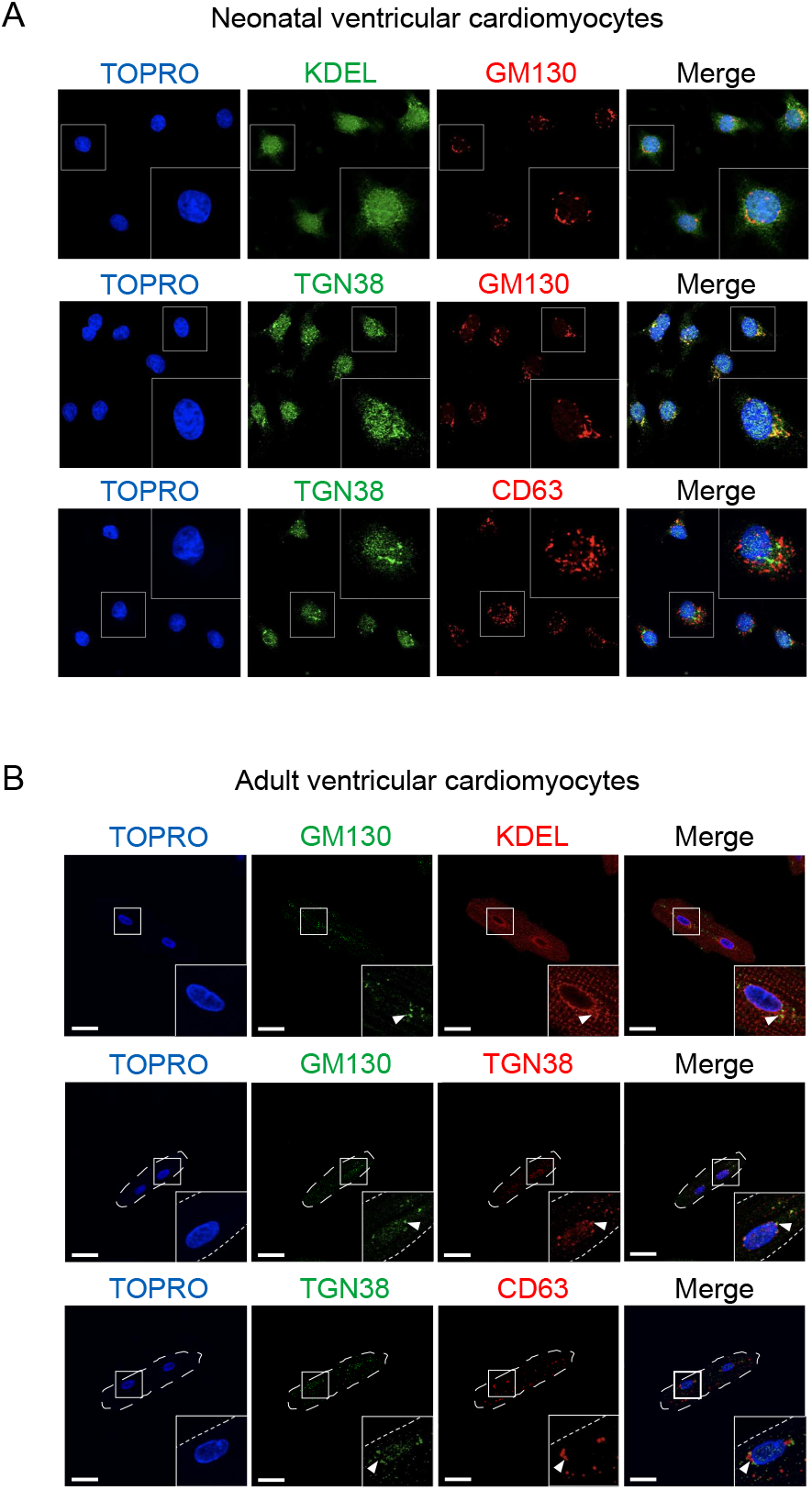
Protein secretion of the classical secretory pathway is predominantly initiated from the ER. **(A)** Coimmunofluorescence labeling of neonatal rat ventricular myocytes with antibodies to the ER-marker KDEL, the cis-Golgi-marker GM130, the trans-Golgi-marker TGN38 and the exosome-marker CD63. **(B)** Coimmunofluorescence labeling of adult rat ventricular myocytes as described in (A). TOPRO, nuclei; bars, 20 µm.

### Ribosomes directly bind to both the sarcoplasmic and endoplasmic reticulum of adult cardiac myocytes

We proceeded to further identify the membrane network associated with protein synthesis in differentiating cardiac myocytes. Accordingly, we performed immunofluorescence co-localization studies of neonatal and adult rat cardiac myocytes for p-RPS6, indicating ribosomes engaged in translation, together with either RyR2, SERCA, or KDEL (Figure 3). Phosphorylation of RPS6 correlates with ribosome activation and translation of selected mRNA transcripts in the heart [19,30-33], which are involved in cell cycle progression, translation and specialized functions to regulate cardiac function [19,30,32,34]. In neonatal rat cardiac myocytes p-RPS6 was found throughout the cell with intense staining at the peri-nuclear ER (Figure 3*A*). In contrast, in adult rat cardiac myocytes p-RPS6 localized to the ER and SR (Figure 3*B*). Additionally, p-RPS6 was localized to the distal ends of the cardiac myocytes near the intercalated disc, which was described previously [16,35]. RPS6 is is a component of the 40S ribosomal subunit [36]. To validate our results we isolated adult ventricular myocytes from RiboTag mice, which we previously used to characterize the protein synthesis response of cardiac myocytes in response to stress [19,37]. RiboTag mice express hemagglutinin (HA)-tagged ribosomal protein RPL22, a subunit of the 60S ribosomal subunit that is incorporated into actively translating polyribosomes [37-39], under a cell-type specific promotor, in our case specifically in cardiac myocytes. Similar to p-RPS6, HA-RPL22 located to both the ER and SR in adult ventricular myocytes, confirming that our staining pattern for ER and SR-localized ribosomes is independent of ribosomal subunit, antibody and species (Figure 3C). If ribosomes attached to the ER and SR are actively involved in translation, mRNAs should in general co-localize to similar subcellular regions. Thus, we used mRNA fluorescence in situ hybridization (FISH) to visualize the distribution of poly(A)-mRNAs in cardiac myocytes (Figure 3D). Mapping of mRNA localization using a probe for poly(A)-mRNAs showed that similar to ribosomes, mRNAs localize to a perinuclear ER-like region and a peripheral striated SR-like network in adult cardiac myocytes (Figure 3D).

**Figure 3.**
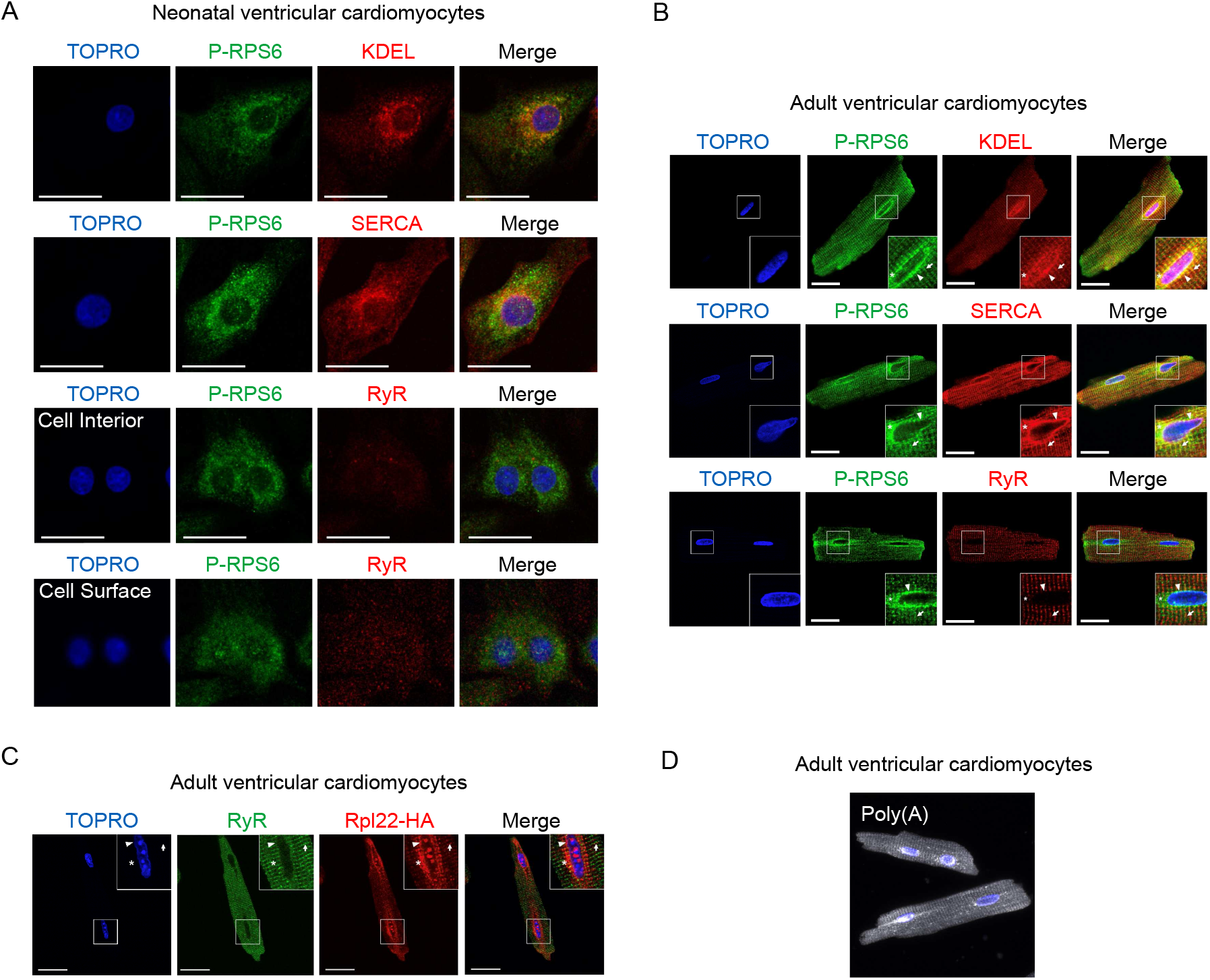
Ribosomes are localized throughout the neonatal and adult ventricular myocyte ER and SR membrane network. **(A)** Coimmunofluorescence labeling of neonatal rat ventricular myocytes or **(B)** adult rat ventricular myocytes with antibodies to the S6 ribosomal protein phosphorylated at Ser235/236 (p-RPS6) and either an antibody that specifically recognizes proteins containing the KDEL sequence, used as an ER marker, the Ca^2+^-ATPase of the SR (SERCA), or the cardiac ryanodine receptor (RyR2). **(C)** Coimmunofluorescence labeling of isolated adult ventricular myocytes from RiboTag mice, expressing hemagglutinin (HA)-tagged ribosomal protein RPL22, which is incorporated into actively translating polyribosomes. TOPRO, nuclei; bars, 20 µm. **(D)** mRNA fluorescence in situ hybridization (FISH) of poly(A)-mRNAs in adult rat cardiac myocytes.

To get further information on the areas of protein synthesis in cardiac myocytes we additionaly mapped the sites of active translation by acute puromycin pulsing. Puromycin is an aminoacyl-transfer RNA structural analogue which incorporates into elongating peptide chains and can be detected by an anti-puromycin antibody. Neontatal and adult cardiac myocytes were treated with 0.5 µg/ml puromycin and rapidly fixed to avoid dissociation of the puromycin-labeled polypeptide [40]. Acute puromycin pulsing showed that protein synthesis localizes to areas of the ER, SR, and the cytosol of neonatal and adult cardiac myocytes (Figure 4A and 4B). This observation is in accordance with other recent reports that describe that cardiac myocytes perform localized translation of selected mRNAs at specific sub-regions [16-18,35] and we postulate that the SR is one such region at which translation occurs in cardiac myocytes. The specificity of the signal was confirmed by co-treatment with the translation inhibitor cycloheximide, which resulted in dissapearence of the puromycin signal.

**Figure 4.**
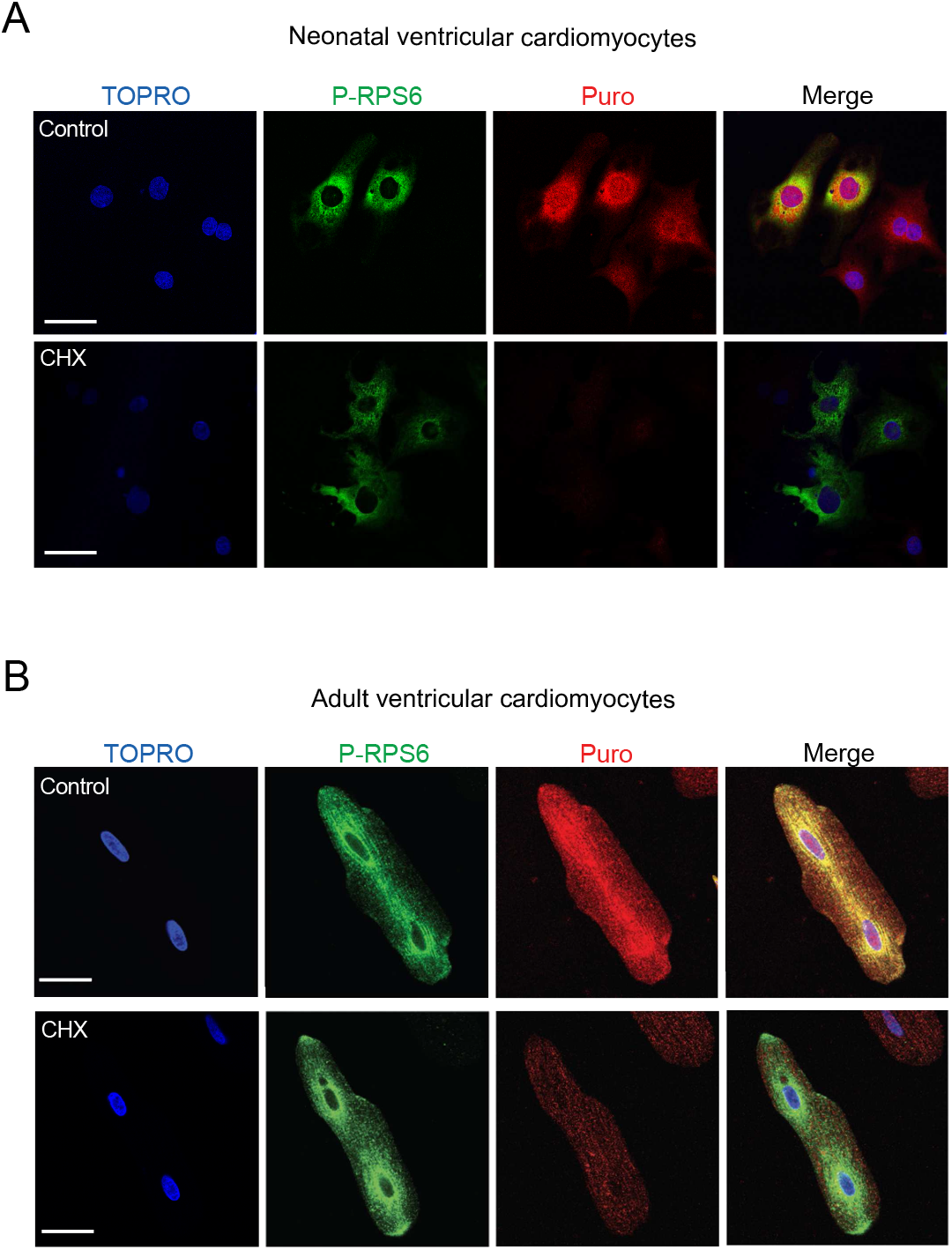
Protein synthesis is localized near the SR and ER in cardiac myocytes. **(A)** Coimmunofluorescence labeling of neonatal rat ventricular myocytes or **(B)** adult rat ventricular myocytes with antibodies to the S6 ribosomal protein phosphorylated at Ser235/236 (p-RPS6) and an antibody that specifically recognizes puromycin incorporated into protein (Puro). Scale bars: 50 µm (A), 20 µm (B). **(C)** Schematic diagram of the sarco/endoplasmic reticulum network in a cardiac myocyte, depicting the relationships between the SR [A] the peri-nuclear ER [B] and nuclear envelope [C]. The localization of secreted and membrane protein synthesis to the nuclear envelope, peri-nuclear ER, and the SR, depicted as a contiguous membranous system, is shaded purple.

### Superresolution microscopy confirms the direct interaction of ribosomes with the sarcoplasmic reticulum in cardiac myocytes

To further interrogate details of ribosome localization to the SR of adult cardiac moycytes, we proceeded with high resolution mapping of the distribution of ribosomes attached to the ER and SR. We used transmission electron microscopy to examine the intracellular relation of ribosomes to membrane networks across different subcellular localizations. Transmission electron microscopy of cryo-prepared adult rat cardiac myocytes showed that membrane-bound ribosomes can be found at peri-nuclear membranes (ER) (Figure 5A), peripheral membrane networks (SR) (Figure 5B), near the t-tubules (Figure 5C), and in subsarcolemmal regions, for example, at the intercalated disc (Figure 5D), confirming our initial observations made by confocal microscopy.

**Figure 5.**
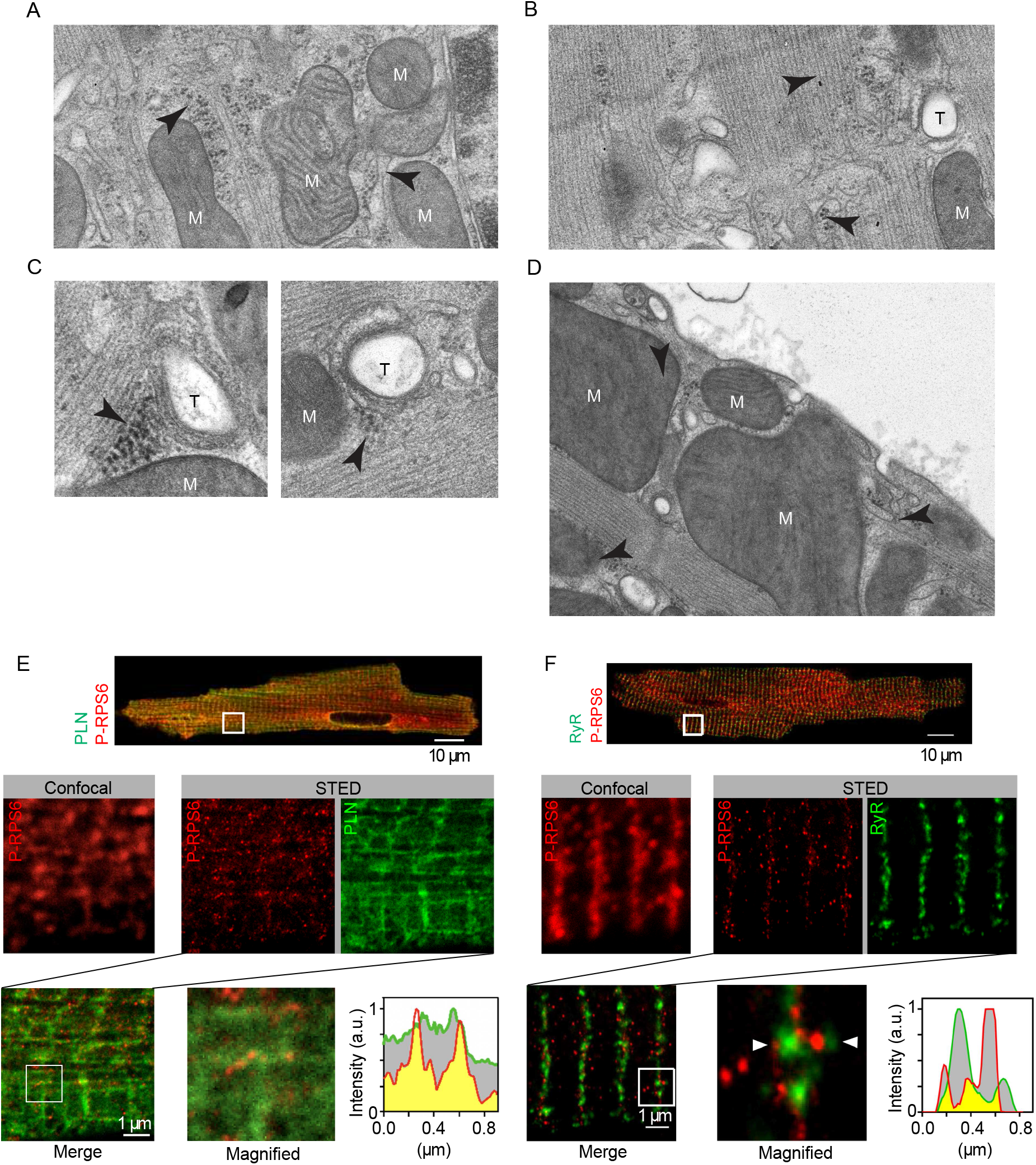
Ribosomes colocalize with the SR. **(A-D)** Representative electron micrographs of adult rat ventricular myocytes show (A) ribosome (arrowheads) association with perinuclear membranes (ER), (B) peripheral membranes (longitudinal, free SR), (C) t-tubules (near junctional SR), and (D) at the subsarcolemmal region, specifically at the intercalated disc. M, mitochondria; T, t-tubules; N, nucleus. **(E)** Coimmunofluorescence imaging of phosphorylated ribosomal protein S6 (p-RPS6; red) and phospholamban (PLN; green) or **(F)** ryanodine receptor type-2 (RyR2) in isolated mouse ventricular myocytes using both confocal microscopy and STED nanoscopy. Zoom-in demonstrating the local association of p-RPS6 with individual PLN or RyR2. White rectangles indicate the orientation of the signal intensity profiling. Colocalized signals are indicated in yellow. White boxes in A and B indicate magnified regions. Scale bars: 10 µm (cell overview), 1 µm (confocal, STED), and 200 nm (magnified view). STED, stimulated emission depletion.

While electron microscopy revealed a clear association of ribosomes with membrane systems across the cardiac myocyte cell body, it did not functionally distinguish different membrane networks, e.g. ER from SR, to clearly confirm that ribosomes are directly attached to the SR and not simply in close proximity to it, as our co-localication imaging by confocal microscopy does not necessarily indicate direct interaction due to its limited resolution. Superresolution microscopy using stimulated emission depletion (STED) with physical resolution improvement is fundamentally diffraction-unlimited and has a resolution of down to 20 nm, which is smaller than the size of a eucaryotic ribosome [36,41]. To investigate whether ribosomes directly interact with the SR, we co-stained adult cardiac myocytes with P-RPS6 (ribosome marker) and phospholamban (PLN) or RyR2 (both classical markers of the SR) using STED microscopy. While the co-staining of P-RPS6 with PLN indicated some colocalization, but remained inconclusive due to the imperfect PLN antibody signal, co-staining of P-RPS6 with RyR2 clearly revealed that ribosomes directly localize to the SR in cardiac myocytes, marked by RyR2 (Figure 5E and 5F). In summary, these findings confirm the interaction of the SR membrane network of adult cardiac myocytes with active ribosomes.

Taken together, distinct but interconnected membrane networks that are involved in functions classically associated with the ER or SR develop and coexist in mature adult cardiac myocytes. Both networks conduct distinct but overlapping functions as indicated by the localization of specific proteins to different regions of the cardiac myocyte membrane network, even though some proteins such as SERCA are found throughout the network. In general, a peri-nuclear membrane system that appears interconnected to the nuclear envelope conducts functions classically associated with the ER of other cell types, such as protein secretion through the classical secretory pathway, while a peripheral network conducts functions classically associated with the SR, such as calcium-induced calcium release. Using a combination of different imaging techniques, including superresolution microscopy, we reveal that ribosomes bind to both the ER as well as the developed SR in adult cardiac myocytes, suggesting that both the ER and SR are sites of protein synthesis.

## Discussion

The ER is a major site of protein synthesis and is involved in the synthesis, folding, modification, and transport of proteins that are critical for the function of excitable cells, such as ion channels, receptors, calcium-handling proteins or secreted proteins. In striated muscle cells, a membrane network functionally different from the ER of other cell types, known as the SR, has been defined and surrounds the muscular myofilaments and controls the release of calcium into the cytoplasm to regulate contraction. However, the relative locations, protein synthetic functions, and protein expression profiles of the ER and the SR in cardiac myocytes are unknown [12]. Using a combination of different imaging techiques, including superresolution microscopy, we show that adult cardiac myocytes contain two functionally distinct membrane networks, a peri-nuclear ‘ER‘ and a peripheral ‘SR‘, which appear interconnected, as shown for skeletal muscle cells [8,12,42].

Cardiac contraction is initiated by the release of Ca^2+^ from the SR in response to an action potential, in a process known as excitation-contraction (EC) coupling. The components of EC coupling are under tight spatial and temporal control and mature during development [43-49]. Postnatal development of cardiac myocytes is associated with reorganization of the cellular architecture supporting an increase in mitochondrial and myofibrillar mass and SR maturation [50]. The main energy consumers of the cardiac myocyte are involved in EC coupling and are localized in the SR and myofibrillar compartments, while energy production is mainly performed by the mitochondria. Therefore, maturation of cardiac myocyte architecture underlies the development of intracellular energy pathways and efficient EC coupling. In line with this, we found that the SR compartment increases from neonatal to adult cardiac myocytes. In adult cardiac myocytes, the SR spreads throughout most of the cellular periphery, while the ER is found exclusively in a smaller peri-nuclear region, likely representive of the cellular need for efficienct Ca^2+^ handling and conctracile regulation.

Our imaging data suggests that secretion of the classical secretory pathway arises primarily from the peri-nuclear ER, similar to other cell types. This observations stands in contrast with findings from Bogdanov et al. who proposed that the machinery necessary for membrane protein synthesis and secretion is distributed throughout the cardiac myocyte [35]. While our imaging data showed a higher density of the secretory machinery peri-nuclear, we also found those elements to a lesser degree in the cell periphery. Differences between both studies may arise from antibodies used or differences in other experimental conditions, however it seems reasonable that secretion may be initiated from both the ER as well as the SR, with a higher specialization of the ER in protein secretion.

In general, two different models for the relationship between the ER and SR of cardiac myocytes and their protein synthethic functions have been proposed [12], based largely on studies in skeletal muscle cells. In a first model, protein synthesis and secretion occurs only at the nuclear envelope and peri-nuclear ER, defining a membrane network primarily involved in protein processing, functionally different from a ‘smooth’ SR, which does not participate in protein synthesis and is specialized in calcium regulation. In a second model, protein synthesis, folding, modification, and transport is localized to a contiguous membrane network of cardiac myocytes consisting both of a peri-nuclear ‘ER-like‘ system and a peripheral ‘rough SR‘. Our data points towards the second model, however, other functions do not appear to be shared between both systems, such as RyR2-mediated Ca^2+^ release. Recently, other studies demonstrated that cardiac myocytes distribute specific mRNAs to different sub-cellular regions where they can be found in close association with ribosomes, indicative of localized translation [16-18,35]. This process appears to depend at least partly on microtubule-associated transport [17]. While it was previously known that the intracellular membrane system of cardiac myocytes is essential for translation and processing of many important proteins such as secreted and membrane proteins, e.g. ion channels or cell surface receptors, differences between the ER and SR and whether the SR participates in protein synthesis and processing remained unclear [12]. Our study adds up on the previous findings of localiced translation in cardiac myocytes by defining the relative locations and differences between the ER and SR of cardiac myoyctes and by systematically analyzing the association of ribosomes to the SR, indicating that the SR of adult cardiac myocytes represents a novel sub-cellular region of localized translation. Polarized cells such as neurons or glia cells synthesize proteins near the site of their final function and this may hold true for ER- and SR-associated proteins. Many Ca^2+^ handling proteins reside in SR, making it reasonable that those proteins are synthesized on ribosomes associated with the SR in cardiac myocytes, similar to the way that ER-luminal and transmembrane proteins in non-cardiac myocytes are synthesized on ribosomes associated with the ER.

In summary, cardiac myocytes contain both an ER and an SR that perform overlapping but distinct cellular functions. The membrane compartment present in neonatal cardiac myocytes resembles mostly the ER and decreases in relative abundance during cardiac myocyte maturation. In the adult cardiac myocytes, the SR is the main membrane network. Both networks can be distinguished by RyR2, which is not present in the peri-nuclear ER but is expressed in the peripheral SR. While protein secretion via the classical secretory pathway is predominantly initiated from the ER, other functions such as SERCA-mediated Ca^2+^ reuptake are shared between both membrane systems. Specifically, using a combination of different imaging methods, including super resolution microscopy, we reveal that similar to the ER, the SR of adult cardiac myocytes binds active ribosomes and is a site of translation.

## Acknowledgments

Ch.S. acknowledges the German Cardiac Society. Ch.S, M.V and S.D acknowledge the German Centre for Cardiovascular Research-Partner Site Heidelberg/Mannheim.

